# Commentary: BRAIN NETWORKS. Correlated gene expression supports synchronous activity in brain networks. S*cience* 348, 1241-4

**DOI:** 10.1101/079202

**Authors:** Spiro P. Pantazatos, Xinyi Li

**Affiliations:** Departments of Psychiatry, Columbia University, New York, NY; Biomedical Informatics, Columbia University, New York, NY

## Abstract

A recent report claims that functional brain networks defined with resting-state functional magnetic resonance imaging (fMRI) can be recapitulated with correlated gene expression (i.e. high within-network tissue-tissue “strength fraction”, SF) (Richiardi et al., 2015). However, the authors do not adequately control for spatial proximity. We replicated their main analysis, performed a more effective adjustment for spatial proximity, and tested whether “null networks” (i.e. clusters with center coordinates randomly placed throughout cortex) also exhibit high SF. Removing proximal tissue-tissue correlations by Euclidean distance, as opposed to removing correlations within arbitrary tissue labels as in (Richiardi et al., 2015), reduces within-network SF to no greater than null. Moreover, randomly placed clusters also have significantly high SF, indicating that high within-network SF is entirely attributable to proximity and is unrelated to functional brain networks defined by resting-state fMRI. We discuss why additional validations in the original article are invalid and/or misleading and suggest future directions.

## Main Text

A recent study explores relationships between gene expression and distributed spatial patterns of synchronous brain activity consistently observed in resting state (RS) fMRI (Richiardi et al., 2015) using microarray data from the Allen Brain Atlas (http://human.brain-map.org, (Hawrylycz et al., 2012)). The authors correctly state that “While functional networks are distributed spatially, meaning they cross over different tissue types, and that their sample can be spatially distant, it is important to ensure that a high strength fraction (SF) does not simply reflect the fact that tissues are the same.” They attempt to correct for spatial proximity by omitting edges between regions falling in the same “tissue class”, which are ontological labels provided by Allen Brain Atlas (Supplementary Table 4 in (Richiardi et al., 2015). However, this approach inadequately controls for spatial proximity: nearby regions will fail to have their edges removed by a label boundary dividing them, while longer edges within a tissue label will be removed instead (Fig 1A). The issues remains even when correction uses coarser tissue classes.

**Fig. 1:**
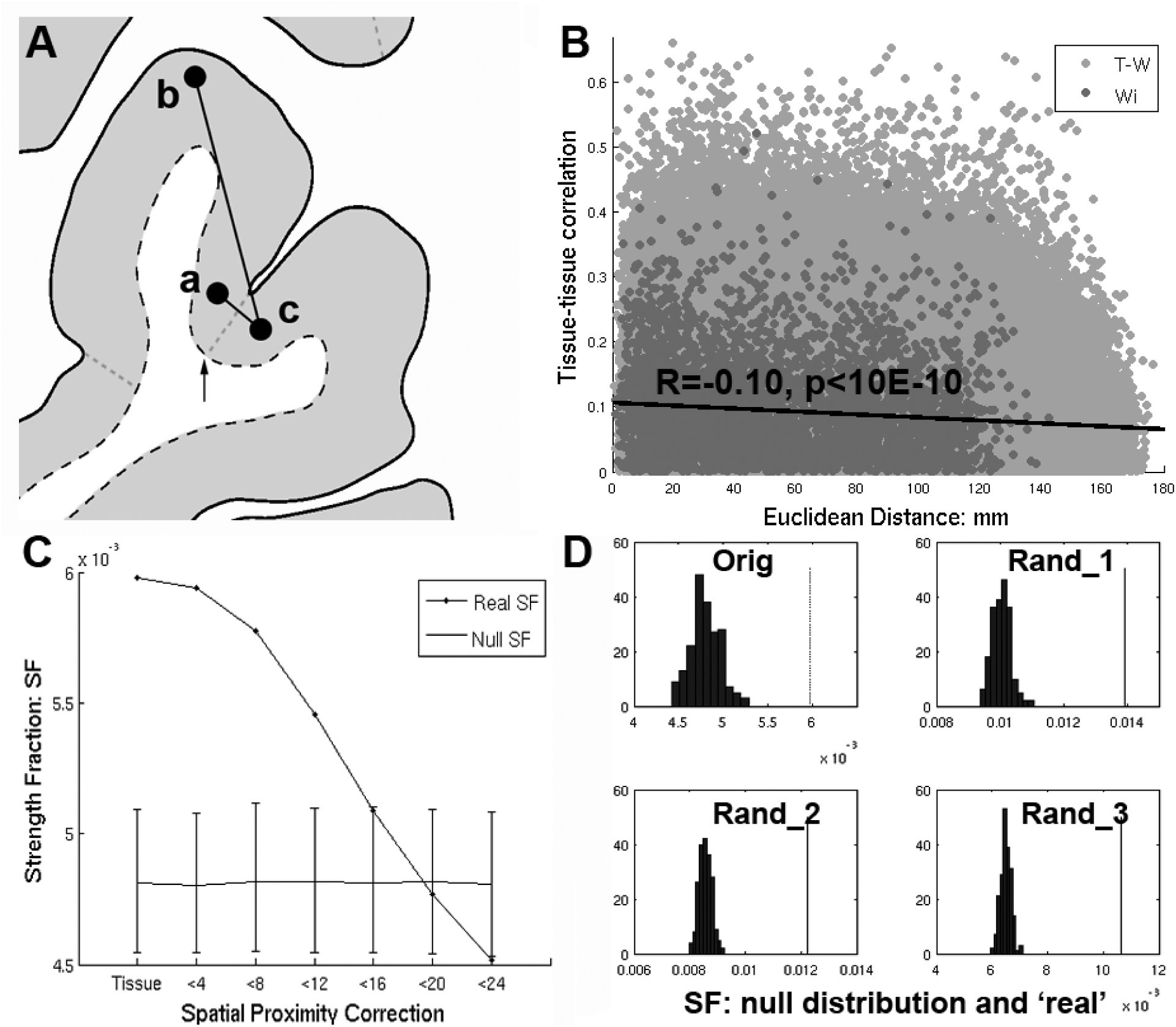
**A)** Richiardi et. al attempts to control for spatial proximity by removing edges with nodes having the same tissue label (i.e. **a** and **b**). However. nearby regions **a** and **c** will fail to have their edges removed by an arbitrary label boundary (arrow) that divides them, while more distant edges (**a-b**) within a tissue label will be removed instead. **B)** Even after removing within-tissue edges, there remains a strong dependence of tissue-tissue correlations on distance (R=−0.10, p<10E−6), with nearby regions tending to have higher tissue-tissue correlations. **C)** Strength fraction (SF) depends on spatial proximity. “Tissue” refers to the original within-tissue class correction applied by Richiardi. et. al. and corresponds to their primary findings (p<10E-4). However as short distances (edges) are removed (< 4 through 24 mm) the SF falls monotonically until it is no longer greater than the null distribution at <20 mm. **D)** Upper left corner (“orig”) shows the null distribution and SF corresponding to the main results reported in (Richiardi et al., 2015), while the rest of the panels show the same for 3 randomly placed sets of contiguous clusters.

Even after removing within-tissue edges, there remains an association between tissue-tissue correlations and distance (R=−0.10, p<10E−6), with nearby regions tending to have higher correlations (Fig 1B). Within network (Wi) edges are significantly shorter than out-of-network (T-W) edges (Wi distances vs. T-W distances 2-sample t-test: t_(759,091)=_−51.1, Wi mu=52.9 mm, T-W mu=78.3 mm). This biases the Wi SF to be greater relative to a null distribution which calculates Wi using longer connections (i.e. T-W edges which are labeled Wi as part of the shuffling).

When a more direct correction for distance is applied (removing proximal edges), within-network SF is no longer greater than null. Fig 1C shows dependence of SF on proximity. “Tissue” refers to the within-tissue class correction applied by Richiardi et al. and demonstrates their primary findings (p<10E–4). However, as short-range edges are removed (4–24 mm), SF falls monotonically until it is no greater than null at <20 mm. In addition, applying linear regression to adjust for distance (French and Pavlidis, 2011) results in a large *negative* SF (SF=−0.61, p=1, data not shown). *Thus, the claim of the original article: “Given that we used only cortical samples, that we removed edges linking tissues of the same class, and that functional networks are spatially distributed, this finding cannot emerge from spatial proximity or gross tissue similarity” is false.* Moreover, the null distribution derived in Richiardi et al. is flawed because the permutation strategy assumes all regions are independent and equally exchangeable, which is not true given the spatial autocorrelation and distance bias.

Although not reported in the original article, the authors claim that SF remains significant after a linear regression-based distance correction is applied and only positive connections are included (personal communication). However, there are two problems with this: 1) The assumption that tissue-tissue correlation strength various linearly with distance is too strong. A plot of the tissue-tissue correlations vs. distance shows that the best-fit curve is steep for short edges and less steep at around 20mm: after adjusting for the best-fit line there will still be a distance bias. Model-based correction will not be as optimal as simply removing proximal connections. 2) Applying a cutoff of zero for connections contributing to the SF is not well justified (this applies to the main analyses as well). What, biologically, distinguishes a correlation of 0.1 vs −0.1 other than i.e. noise in the expression vector? Furthermore, after regression, about half of the connections (that were included in the original main analyses) will be negative due to mean centering and omitted in the new analysis, making the cutoff of zero even more arbitrary.

Short (<16 mm) connections account entirely for the significant SF reported in (Richiardi et al., 2015) (Fig 1C). Given that the main claim of Richiardi et al. is that correlated gene expression relates specifically to RS functional networks, a crucial question is “is high local SF specific to the RS networks”? If so, then the SF of distributed clusters with centers randomly placed throughout cortex (with size and total number of Wi nodes similar to RS networks) should be non-significant. However, for 1,000 randomly selected cluster sets, p-values were *all 0 (<0.001).* Panel D (“Orig”, upper left corner) shows the null distribution and real SF corresponding to the main results reported in the original article, while the rest of the panels show the same for 3 randomly selected sets of clusters. Thus the significant SF reported in the original article is entirely attributable to spatial proximity *and* is unrelated to RS fMRI networks. Note that SF of Wi RS networks cannot be compared to SF of randomly selected clusters since SF is a function of total number of Wi connections.

Matlab (and eventually R) code replicating the primary results presented in (Richiardi et al., 2015) and results presented here are available at https://github.com/spiropan/ABA_functional_networks. See Supplement for a discussion why the additional validation analyses in (Richiardi et al., 2015) (Figures 2 and 3) are invalid and/or misleading, and the relationship of their results with differentially stable genes identified in (Hawrylycz et al., 2015).

## Acknowledgments

This work was supported in part by a Janssen Translational Neuroscience Postdoctoral Research Fellowship and a K01MH108721 (SPP). We thank Paul Pavlidis and Leon French for helpful comments and suggestions.

